# Cell Typing and Sub-typing Based on Detecting Characteristic Subspaces of Morphological Features Derived from Neuron Images

**DOI:** 10.1101/2023.09.26.559442

**Authors:** Sujun Zhao, Penghao Qian, Lijuan Liu

## Abstract

**Motivation:** Recent advances in reconstructing 3D neuron morphologies at the whole brain level offer exciting opportunities to study single cell genotyping and phenotyping. However, it remains challenging to define cell types and sub-types properly.

**Results:** As morphological feature spaces are often too complicated to classify neurons, we introduce a method to detect the optimal subspace of features so that neurons can be well clustered. We have applied this method to one of the largest curated databases of morphological reconstructions that contains more than 9,400 mouse neurons of 19 cell types. Our method is able to detect the distinctive feature subspaces for each cell type. Our approach also outperforms prevailing cell typing approaches in terms of its ability to identify key morphological indicators for each neuron type and separate superclasses of these neuron types. the subclasses of neuronal types could supply information for brain connectivity and modeling, also promote other analysis including feature spaces.

**Availability:** All datasets used in this study are publicly available. All analyses were conducted with python package Scikitlearn 0.23.1 version. Source code used for data processing, analysis and figure generation is available as an open-source Python package, on https://github.com/SEU-ALLEN-codebase/ManifoldAnalysis

## 1 Introduction

The problem of classifying neurons has been studied both qualitatively and quantitatively (Cajal and Azoulay,1952), but falls into disagreements in lumping-splitting cell types (Zeng, 2022). Molecular and anatomical approaches, including the profiling of RNA transcripts (transcriptomics) as well as characterization of spatial distribution and morphology of single neuron, are prevalent in cell type classification (Zeng, 2022; Peng et al., 2021; Gouwens et al., 2019). It is common to relate transcriptomic profiles with other modalities (Gala et al., 2019) since effective single-cell molecular profiling can be easily achieved with traditional clustering approaches (Figure 1 A2). However, the extent to which transcriptome clusters can represent true cell types and what level of granularity is appropriate is still a question. For definitive cell type classification, a full representative set of morphology and connectivity, which are regarded as the most defining features for neurons since the era of Cajal, is required (Zeng, 2022). Neural morphology and connectivity form the physical system of the brain (‘hardware’ base) (Pfeifer and Gomez, 2009), so they are pivotal to understand how a brain works. In fact, neuron morphology alone can be used to identify cell types or subtypes, such as five neuron classes of layer-5 neurons of mouse primary visual cortex determined using hierarchical clustering (Tsiola et al., 2003). Besides morphology is quite robust to experimental conditions among the features commonly used to describe neurons (Ascoli and Wheeler, 2016). But morphology-based cell typing remains unclear considering that morphological characteristics are difficult to comprehensively capture and too complicated to cluster by traditional methods, unlike transcriptomic data.

**Figure 1.**
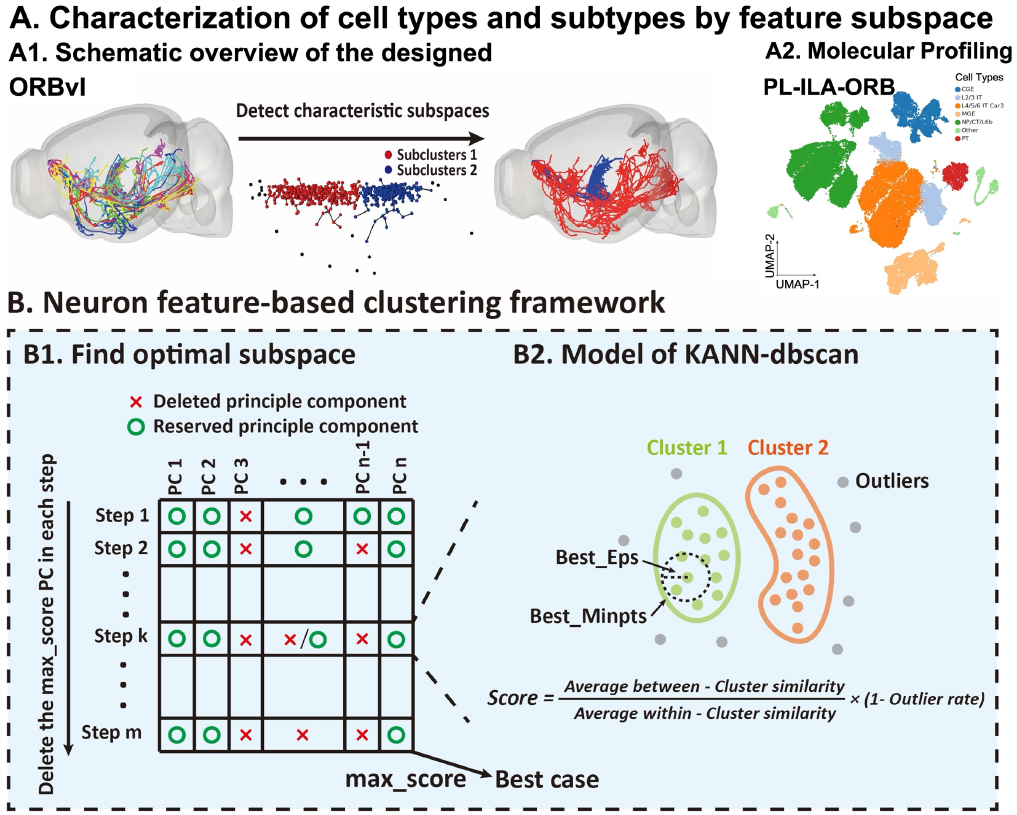
Workflow of the complete feature subspace selection. **(A)** Characterization of cell types and subtypes by feature subspace. **(A1)** A given dataset of neurons can be well classified under the optimal characteristic subspace detected by our algorithm, where neurons show their greatest specificity and intrinsic patterns. **(A2)** Cell typing and subtyping under the detected feature subspace can correspond to the molecular profiling to some extent. **(B)** Comprehensive framework for feature subspace selection and clustering. **(B1)** In order to find the optimal subspace, the algorithm iteratively deletes a feature vector in turn during each dimension reduction step and reserve PCs which can generate the maximum score into the next step, repeating until the dimension reaches 3 or the result cannot be split. **(B2)** KANN-DBSCAN model, able to automatically to adjust parameters, runs with the remaining eigenvectors and then the score((intersimilarity/intra-similarity) × (1-outlier%), see Methods) is computed. Among all the results, the case with the largest score is the best, and the corresponding feature space is the optimal feature subspace.

With the rapidly available brain-wide, digitally reconstructed single neuron morphology from high-resolution microscopy images (Winnubst et al., 2019; Peng et al., 2015), the number of neuron morphology datasets has sprung up from dozens to thousands during the last twenty years. A cutting-edge research field is how to analyze these datasets with sufficient biological relevance. Typical way is to first extract morphological or topological features, like L-measure features that describe global attributes of neuronal structures (Scorcioni et al., 2008), and then choose a clustering algorithm to classify neurons (McGarry, 2010; Karagiannis et al., 2009; Armañanzas and Ascoli, 2015, Peng et al., 2021). Considering the difficulty in validating results from unsupervised learning and the applicability of clustering algorithms, finding an optimal algorithm has always been focus of the study (Masood, M. and Khan, M.N.A., 2015). In contrast, representation and feature learning for clustering have not been explored extensively (Karim et al., 2021).

Many existing feature selection methods screen features subjectively with prior biology knowledge or automatically by dimension reduction algorithms (Marx and Feldmeyer, 2012; DeFelipe et al.,2013; Armañanzas and Ascoli, 2015). This does not necessarily improve the clustering quality but sometimes causes loss of important features instead. Besides, researchers usually increase the number or variety of features, optimize network architectures, or use more complicated classifiers (Kanari et al., 2019; Lin and Zheng, 2019; Deitcher et al., 2017; Gillette and Ascoli, 2015; Wan et al., 2015) to obtain better clustering results. Although these approaches could categorize neurons to some extent, it is difficult to separate neurons with complex morphologies. Differentiating highly similar neuron structures is also a challenge.

This paper introduces a method to identify characterizing feature subspace where neurons may be optimally grouped. We rank the data clustering in a series of feature space subsets according to a separability measure, so that within-class and between-class discrimination are optimized while the integrity of information is retained. Applying our method in a neuron morphology dataset containing over 9,400 mouse neurons of 19 cell types, we have identified morphologically distinct neuronal subtypes from certain anatomical brain regions. Comparison with other classic clustering approaches illustrates the effectiveness and advantages of our method.

## 2 Results

### 2.1 Optimal subspace-based clustering framework

The primary scope of our study is classification of single neuron morphology by automatic detection of the optimal feature subspace. Under the detected feature subspace, retained structure characteristics is consistent with molecular profiling to a certain extent (Figure 1A). We developed a framework where the search strategies employed are greedy sequential searches through subsets of the original feature space, followed by an evaluation for each of them (Blum and Langley, 1997). Like most approaches to subspace analysis, our measurement is based on a density-based clustering notion (Procopiuc et al., 2002; Baumgartner et al., 2004). The screening-retention process is evaluated by a score (see Materials and Methods) computed from clustering results from the density-based algorithm DBSCAN (Density-Based Spatial Clustering of Applications with Noise).

DBSCAN has two major parameters, the neighborhood size, Eps, and the number of data objects in that neighborhood, MinPts (Galán, 2019). And KANN-DBSCAN, a self-adaptive algorithm used here, is capable of automatically acquiring the best combination of the two parameters for DBSCAN. Considering that DBSCAN outputs both clusters and outliers, the customized evaluation score is formulated by both Euclidean distance between data points and the number of outliers (Figure 1 B2), which reflects not only the aggregation within a cluster and separation between clusters, but also the extent of information loss. Larger the score is, better the classification result, which means the optimal feature subspace is the one under which the highest clustering score is achieved.

The complete procedure goes as follows. The initial step involves data preprocessing and orthogonal transformation into an eigenspace with PCA (Principle Component Analysis). For each step afterwards, all subsets with one eigenvector removed from the current feature space are tested. The dataset is clustered with KANN-DBSCAN, a self-adaptive algorithm for DBSCAN, and then the corresponding score is calculated for each subspace. The feature subspace with the highest score will be reserved as input to the next step. One dimension is reduced in one step and such process will repeat iteratively until the dimension reaches 3 (Figure 1 B1).

We tested our method on 19 cell types totally, where ‘cell type’ here is defined by soma locations in the standard mouse brain atlas: ACAd, ACAv, AId, AIv, CP, FRP, ILA, LGd, MOp, MOs, ORBl, ORBm, ORBvl, PL, SSp, SSs, SUB, VPL, VPM (Jones et al., 2009; Dong, 2008). Overall, there are two outcomes: 1. Subdivisible cell types where neurons can be clearly divided into multiple clusters. Based on whether there exists some correspondence between clustering results and their anatomical locations, two more groups are identified (Figure 2A 2B); 2. nonsubdivisible cell types (Figure 2C).

**Figure 2.**
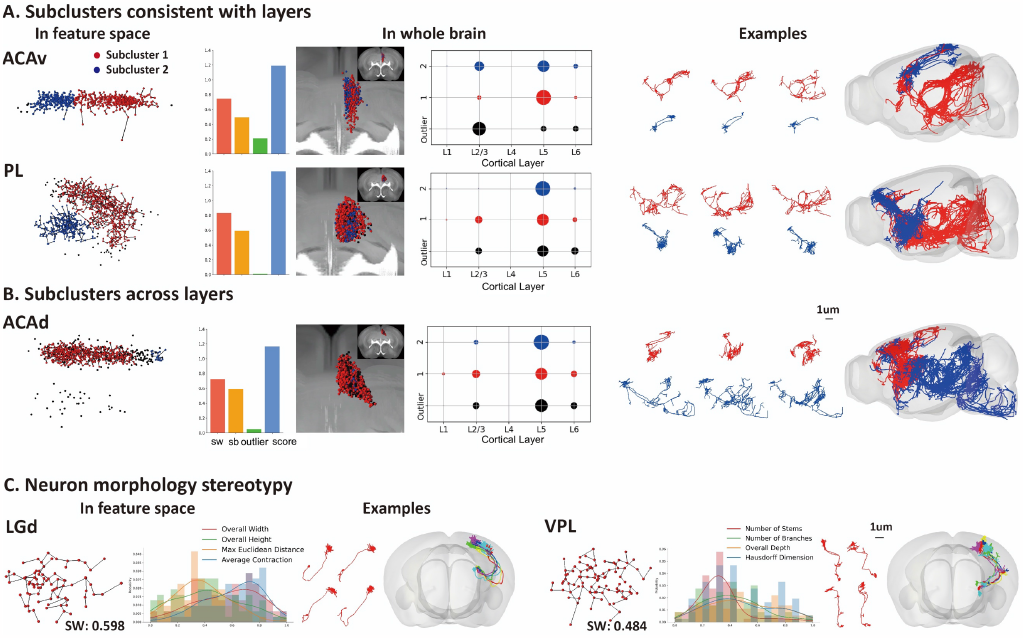
Classification results of single cell types. **(A B).** Sub-divisible cell types. Each point represents a neuron cell. 13 out of 19 types of neurons are considered well separable using our algorithm, exemplified by specific neuron morphologies visualized in the optimal PC subspace and mapped to the CCF brain template (right panel). Sub-clusters from both are distinct to each other in their locations and morphologies, clearly seen in the brain map. There is a correspondence between the division of clusters and soma locations in **A** but not in **B** (middle panel). Due to uniform feature distribution **(C)**, some cell types are considered completely non-divisible here: ILA, LGd, MOp, VPL, SSs and VPM.

### 2.2 Influence of algorithm’s parameters and performance analysis

As we mentioned before, DBSCAN two parameters, Eps and MinPts (Galán, 2019), which have been proved to have a great and import influence on the clustering result (Figure 3A). KANN-DBSCAN can help to find the best combination after automatically trying all parameter pairs. Next, to evaluate the impact of feature subspaces, we visualized the results of all subsets. As the feature subspace changes, data distributions and clustering results are quite different even for the same dimension (Figure 3C). As the dimension is reduced, the score (see Methodology) also varies a lot (Figure 3B). Thus, through adjustment of model parameters and appropriate selection of feature spaces, we can find the optimal feature subspace for a best classification. We have also noted that when separating the superclass dataset (e.g. VPM and CP), the difference in quantity of cell types could affect the final optimal subspace remained and the compactness of clusters (Figure 3D). But the concordance of classification produced three datasets that consist of different components verifies the stability and robustness of our method, and its effectiveness on super-class classifications too.

**Figure 3.**
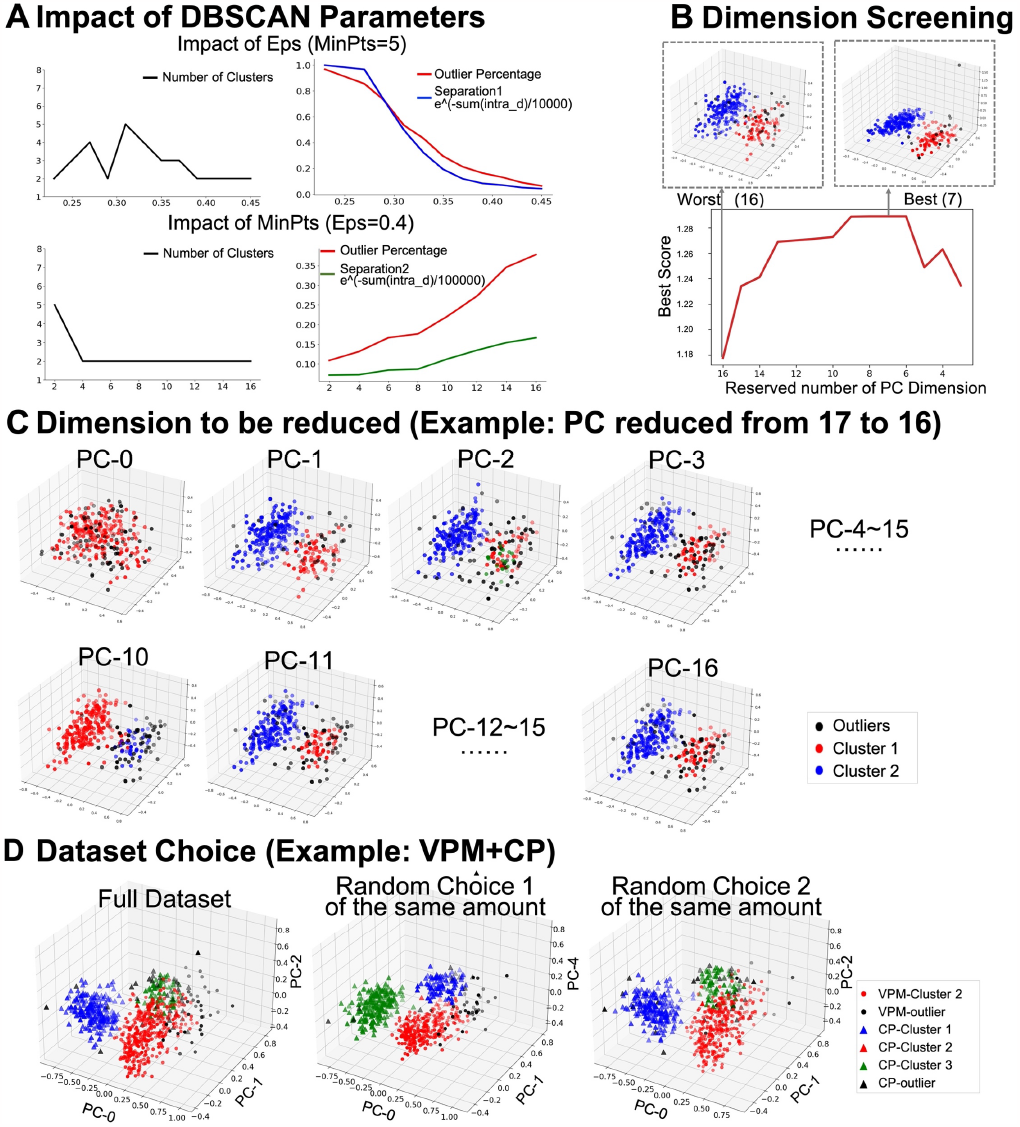
Impacts of algorithm parameters and dataset selection on classification results (CP cells as an example). **(A)** Parameter sensitivity of DBSCAN model. DBSCAN has 2 major tuning parameters: Eps (left) and MinPts (right). Different choices of the two parameters will lead to different classification results. **(B, C)** Influence of feature subspace. **(B)** The score ((inter_dist/intra_dist) × (1-outlier%), see Methodology) varies with the reduction of PC dimensions and the peak corresponds to the optimal dimension of PC subspace. **(C)** The 3D visualization indicates distinct distributions of CP cells in subspaces with the same dimension (16) but different principle components. **(D)** Effect of dataset sampling. Due to the difference in data volumes (e.g. VPM:406; CP: 312), we analyzed the full dataset (718 neurons) and 2 datasets with equal numbered VPM and CP neurons (randomly selected 312 neurons from the VPM dataset). The three results are consistent, indicating the conservation of our algorithm.

Exactly how good is the performance and how much improvement our method would make in morphological classification? For this purpose, we compared our method with 4 typical clustering methods: KMeans, affinity propagation, hierarchical clustering and normal DBSCAN. We used all 5 algorithms to classify datasets of single cell type and conducted morphological analysis on each result. While results were overall comparable, there were indeed some differences between our method and the other four. Taking PL neurons as an example, quantitative comparison of most differentiating morphological indicators shows more prominent distinctions between sub-clusters achieved by DBSCAN than those by the other three (Figure 4A). Besides, DBSCAN preserved underlying continuous manifolds better.

**Figure 4.**
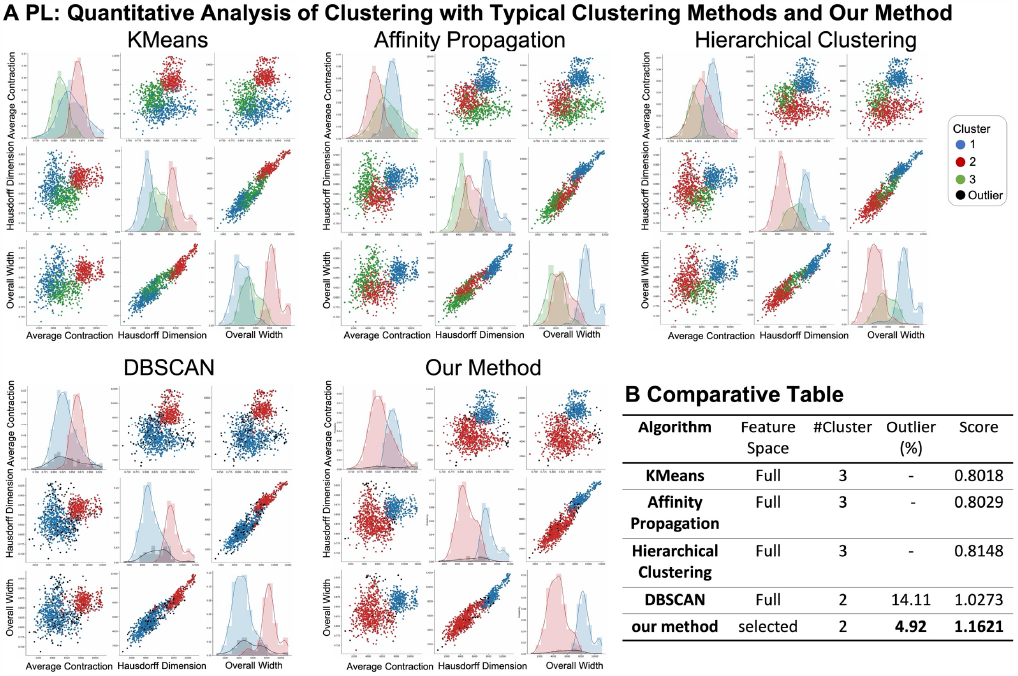
Comparison with typical classification algorithms. 4 typical algorithms are chosen to compare with our method: KMeans, affinity propagation, hierarchical clustering and normal DBSCAN. **(A)** Clustering results of PL neurons, visualized with three most significant morphological indicators: average contraction, Hausdorff dimension and overall width. The subclusters identified with DBSCAN method show obvious differentiations both in morphological structures and under the PC feature space. **(B)** The table verifies that DBSCAN works better than the other three. On this basis, our method turns out to have less outliers and better separation and cohesion (reflecting in the score), compared with DBSCAN without optimization of feature spaces.

Among all the methods, our method has the biggest score, indicating a better classification result. Note that our method (in optimal subspace) performed similar to the normal DBSCAN, but had less outliers, meaning more information was kept (Figure 4B). This emphasizes the role of the optimal feature subspace. Notably, our method is primarily intended to detect the optimal subspace. As above, we could tell features of the highest interestingness were reserved in our method, making it easier to extract the key characteristics for sub-clusters in such feature space (Zhang et al., 2005).

Altogether, the analysis demonstrates the stability and performance improvement of our method when confronted with datasets of different cell types, confirming that the optimal feature subspace may advance the classification of neuron cells and help with the morphological characterization.

### 2.3 Detecting optimal feature subspace and identifying cell subtypes

#### 2.3.1 Identification of key morphological features and multiple subtypes

To gain explanatory power of the optimal feature subspace on its benefits to morphological characterization and categorization, we visualized and quantified the clustering results. In one sub-cluster situation, neurons are morphologically similar to each other. The morphological features have nearly Gaussian distributions in single cell type so that their distribution is homogeneous and sparse under the feature space (Figure 1C). While mostly, subclusters divided actually show significant morphological distinctions from each other. Here, we take ACAv and CP as examples. For CP, 7 eigenvectors are preserved (PC-0,4,5,7,10,16) (Figure 5A). The most significant differential morphological features are mean Euclidean distance, mean path distance, overall width and overall depth (Figure 5B). The two sub-clusters of CP neurons both grow densely at the two ends and smoothly at the long projection part. Neither of them has large branches along the bypassing fibers. The most obvious indicator is the length of the long projection part, clearly seen from the representative neurons, reflecting in the features like Euclidean and path distances (Figure 5C). As for ACAv, the optimal PC subspace has 3 eigenvectors reserved (PC-0,12,14). We checked the correlation between PC subspace and its original features. The main difference lies in overall width, mean Euclidean distance, mean path distance, overall height, overall width, overall depth and total length (Figure 5A 5B) according to the heatmap. Figure 5C validates our result that representative neurons from the two sub-clusters differ remarkedly in branching complexities and overall sizes. These observations and analyses highlight the ability optimal feature subspace to identify sub-type of neurons and the key morphological features.

**Figure 5.**
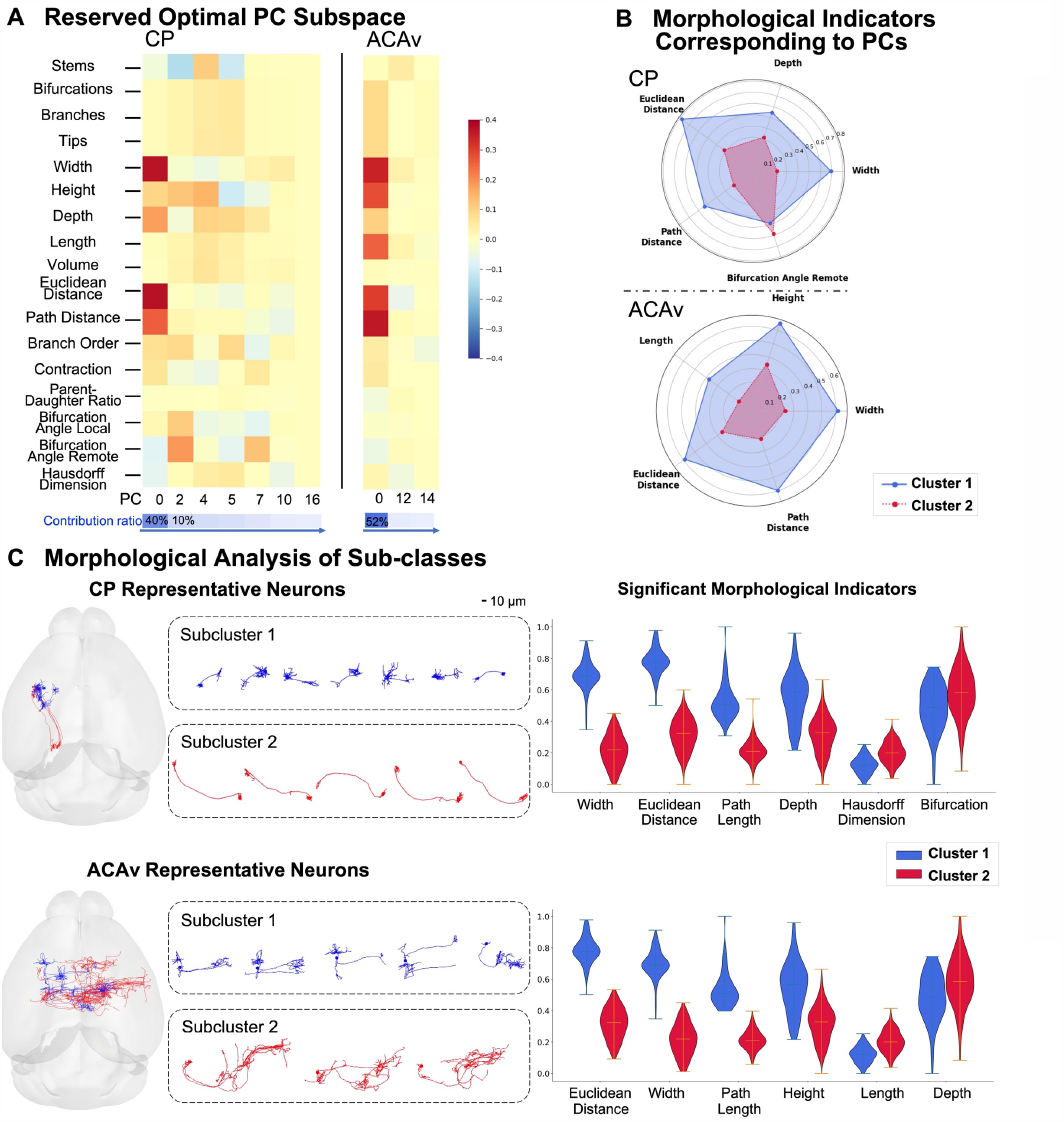
Analysis of clustering results with multiple sub-clusters. Among 13 separable cell types, clustering of 720 ACAv neurons (upper part) and 312 CP neurons (lower part) both have 2 distinct subtypes. (A) Analysis of optimal PC sub-feature space. The heatmap displays the correlation between the optimal reduced components and the original morphological features. Darker the red is, more important the original feature is. And the blue color bar below indicates the contribution ratio of each component, which gets smaller from left to right. (B) According to the heatmap, the most significant morphology indicators are selected with contribution ratio over 0.6. (C) Morphological comparison between two sub-clusters. Left panel: representative neurons for each sub-cluster including their relative locations in the brain and detailed morphological structures. Right panel: distribution of most distinct morphological features (min-max standardized) for each sub-cluster.

#### 2.3.2 Explorative analysis for unsatisfactory clustering

Overall, our method works well on most classification of single cell type but some are exceptions. Due to the morphological specificity of certain cell types, which have uniform and widespread feature distribution, it is hard to conduct a clear subdivision. These kinds of neurons are considered morphologically hard separable. For instance, SSp neurons cannot form large groups where neurons are neither highly similar to each other nor highly different from each other in structures. They distribute dispersively in the optimal subspace, so that the sub-clusters are not so compact and badly differentiated (Figure 6A). On the other hand, sometimes we can only separate neurons based upon their reconstruction sources. In the MOs case, neurons from ION are more easily recognized (Figure 6B). The ION data are structurally incomplete, lack of dendritic parts, making them essentially different in morphologies from data from other two sources, Janelia Mouselight and SEU-Allen. The two sets of data are far apart in the clustering result. Therefore, we clustered these two sets separately to see whether our method still worked. It turns out the sub-clusters obtained are more distanced between and tightly packed within, which proves the robustness of our method on the contrary.

**Figure 6.**
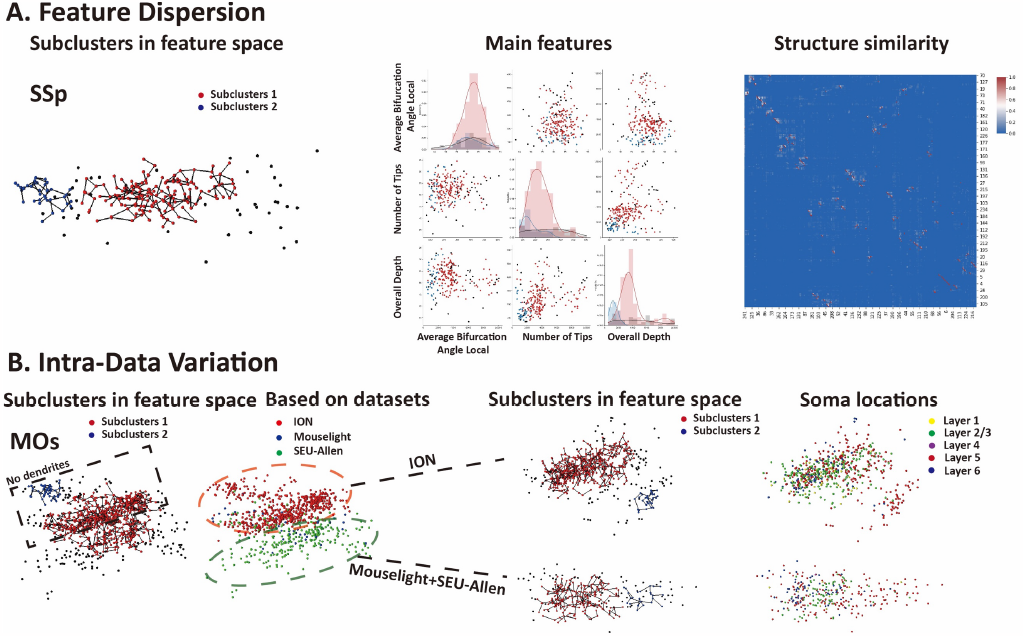
Explorative analysis for unsatisfactory clustering cases. **(A)** Due to feature dispersion. The left panel displays clustering results of SSp and the morphological analysis correspondingly. The distribution of SSp neurons in the feature space is relatively dispersed. Heatmap shows structural similarity between pairs of SSp neuron cells, where the similarity here is calculated by ‘distance’, defined as average distance among all nearest point pairs of neural structures. The neurons are neither highly similar nor highly different to as we can see, so it is intrinsically difficult to achieve an ideal clustering. **(B)** Due to differences in the neuron dataset. Data from ION do not have dendritic parts but make up the most of MOs. These ION neurons are uniformly distributed in the optimal PCA subspace, making them hard to be separated. Therefore, we conducted clustering separately: (1) ION; (2) Mouselight+SEU-ALLEN. The results are much better.

### 2.4 Applications in recognition of multiple cell types from super-class datasets

So far, we mainly focused on the subtyping problem of individual cell type. We conducted this super-class separation to further validate capability and applicability of our method. Since we already have the prior biological knowledge, it is easier to do quantitative check if the classification result is satisfactory or not. As is known that CP neurons have two subtypes (Figure 5), in the first result clearly CP is divided into two groups. And LGd can be considered separated partially at the first place, since LGd neurons seem structurally similar to one sub-type of CP neurons. After the second round of clustering, both LGd and one sub-cluster of CP are identified. The ACAv and VPM case is similar, only that part of ACAv are still mixed up with VPM neurons (Figure 7A). Then we tried a dataset containing three different types from cortex. After two rounds of clustering, the overlapping among cell types does not decrease much. But remarkably, we observed that most AId neurons were separated from the dataset. The third result indicates that roughly cortical neurons share a relatively high structural similarity, among which some types are more recognizable, supporting that there exists some concordance between anatomical regions and neuronal morphology (Figure 7B).

**Figure 7.**
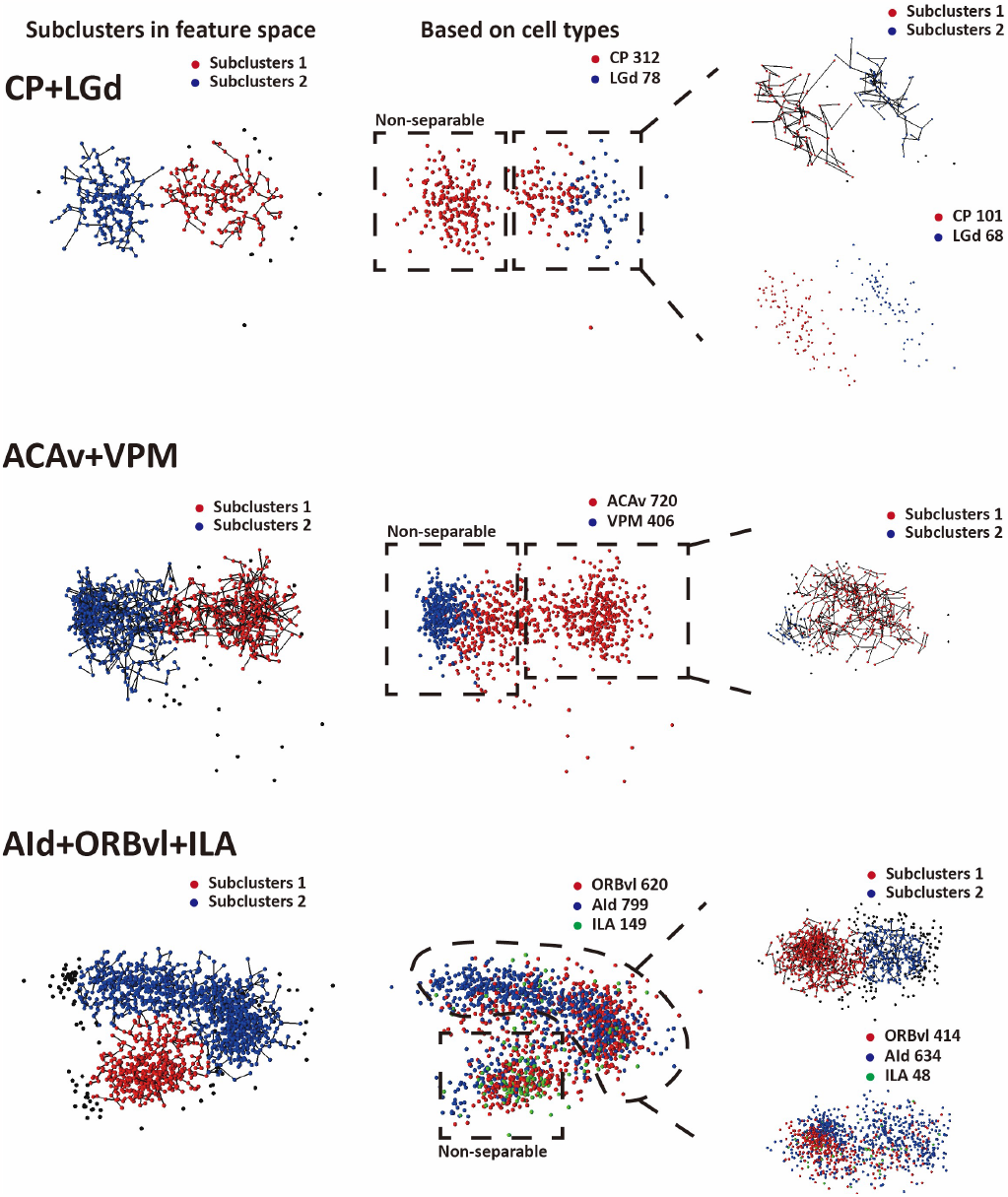
Super-class separation (super-class: ensemble of multiple cell types). 3 datasets are tested: CP(Striatum) and LGd (Thalamus), ACAv (Cortex) and VPM(Thalamus), AId, ORBvl and ILA (Cortex but different cortical layers). Clustering process can be iterated more than one time, and finally cell types are distinguished from the super-class.

In conclusion, with our method, neurons from different brain regions (roughly labeled, e.g. thalamus, cortex) could be separated. While the results also suggest there is some morphological consistency in neurons from the same region. And there is a potential that such super-class problem could be well explored with iteration of exploiting our method.

## 3 Discussion

Previous classifications based on neuron morphology are focused more on dendritic structure mainly, long projection pattern or cross-modality analysis (Milosevic et al., 2009; Cervantes et al., 2019; Peng et al., 2022). In this study, we proposed a new framework for detection of optimal feature subspace and identification of sub-types of neurons. We employed a novel strategy combining greedy optimal eigenspace selection and KANN-DBSCAN to find out such subspace and the corresponding intrinsic morphological features. We used a dataset of 9187 neuron reconstructions from three different sources (ION, Janelia, SEU-ALLEN) and applied our method to decompose single cell types. It turned out 13 of 19 cell types could be separated in the optimal feature subspace we detected. And for the poorly separable cases, most are morphologically similar within cell types.

The whole framework as well as the algorithm were validated in several aspects. First, from the perspective of the algorithm itself, we investigated the influence of model parameters and feature screening on the classification results. Taking these, we optimized the scheme of our method by exploiting the Greedy strategy and introducing a self-defined evaluation score. Second, we found our classification results outperformed those from other algorithms that have been widely used before. Quantitative morphological analysis of identified sub-clusters further confirmed the robustness in that the selected dimensions well preserved significant features, and neurons from different subclusters showed great differences in their structures. Besides, we could distinguish neuron cell types, which are previously labeled, in a super-class group with applications of our algorithm. In general, our method demonstrates great power in detection of key morphological features and classification of neuronal cell types.

However, there are still several limitations. The core idea of our method is to try all the subsets of space dimensions and figure out the best among them, so it will be quite time consuming when the dataset has dozens or even hundreds of features. And as we have discussed in Figure 6A, neurons like SSp with relatively uniform distribution in the feature space are not well classified. Addressing these issues will require future work. Furthermore, some findings in this study deserve more researches. We have discovered some connections between morphological characteristics and soma locations (Figure 2A 2B). The question here is if there exists a pattern, especially for the cortical neurons. For instance, a new subtype from Cortical Layer 5 (L5) has been revealed, showing distinct morphology, physiology and visual response, unlike previously described L5 cortico-cortical and cortico-subcortical types (Kim, E.J. et al., 2015). The cerebral cortex is populated by excitatory and inhibitory neurons of high diversity, which plays a vital role in mediating interactions between various brain regions. But current studies demonstrate mainly on a limited number of types, such as L5 pyramidal neurons. Taking the result we achieved here as the first step, we could be able to build a more detailed model in morphological structures of cortical neurons across layers.

Since we have proved its feasibility in dividing super-class dataset, it is possible to identify an unknown neuron based on its morphological structures, like telling where the neuron locates, through establishing models in optimal feature subspace. However, the question that needs to be answered is that when should we stop the iteration of our algorithm, which is when the clustering is suitable to distinguish both cell types and their subtypes or if these types are truly have significant differences.

In summary, our study highlights the value of detecting optimal feature subspace in classification of neural cells. With the classification and characterization results based on our method, we could establish a standard for morphology-based cell type division. And it is possible to bridge the neural morphology with other modalities such as anatomical definition, electrophysiology and so on, so that if the morphology of an unknown neuron is given, we can tell its cell type and other related biological information. As the number of single full neurons increases, the subtype analysis will be more accurate and better for separation of more cell types. Also, our method provides clues for brain connectivity and modeling and the complexed subtypes in brain promote the other feature space analysis and artificial intelligence. The updated method in neuronal analysis will help more other fields.

## 4 Materials and Methods

### 4.1 Dataset

We used totally 9187 complete morphologically reconstructed neuron structures in the standard SWC file format (Cannon et al., 1998), of which 1741 from SEU-ALLEN, 1200 form MouseLight (Winnubst, J. et al., 2019) and 6246 from ION (Gao, 2022), involving 173 anatomical regions defined from the Allen Reference Atlas ontology. Among them, the ION data does not have dendritic parts. All these data have been registered to the CCFv3 framework. Then we focused on 19 cell types (based on the anatomical regions of somas) with the highest amount and most interest for analysis.

### 4.2 Feature extraction and pre-processing

23 morphological features were extracted for each neuron with the plugin ‘global_neuron_feature’ from Vaa3D platform (Scorcioni et al., 2008) and we kept 17 of them which can better describe their morphologies: “Number of Stems”, “Number of Bifurcations”, “Number of Branches”, “Number of Tips”, “Overall Width”, “Overall Height”, “Overall Depth”, “Total Length”, “Total Volume”, “Max Euclidean Distance”, “Max Path Distance”, “Max Branch Order”, “Average Contraction”, “Average Parent-daughter Ratio”, “Average Bifurcation Angle Local”, “Average Bifurcation Angle Remote” and “Hausdorff Dimension”. These features then are normalized to 0-1 with a min-max standardization and transformed with PCA into a new PC space.

### 4.3 KANN-DBSCAN

DBSCAN is typical of density-based spatial clustering algorithms. The prominent advantages include: 1. no need to specify the number of clusters in advance; 2. outliers can be labeled while clustering; 3. dense datasets of any arbitrary shapes can be clustered. As is known that DBSCAN is very sensitive to its parameters Eps and MinPts. Therefore, in order to achieve the best clustering performance, it is necessary to search for the best combination of the two. We chose KANN-DBSCAN (Li et al., 2019), a self-adaptive algorithm that can automatically determine the parameters. This algorithm, based on the parameter optimization strategy, first generate candidate Eps according to the dataset’s own distribution properties and candidate MinPts. Then it runs DBSCAN repeatedly with these possible parameters to find the stable interval for cluster number change among all the clustering results. The optimal pair of parameters is the one with the minimum density threshold in this interval.

### 4.4 Performance assessment

The result of DBSCAN model includes outliers (invalid points) and clustered points. Referencing Silhouette Coefficient (Kaufman et al., 2005), we designed a score to evaluate its performance, formulating by the data retention rate and the difference of clusters.

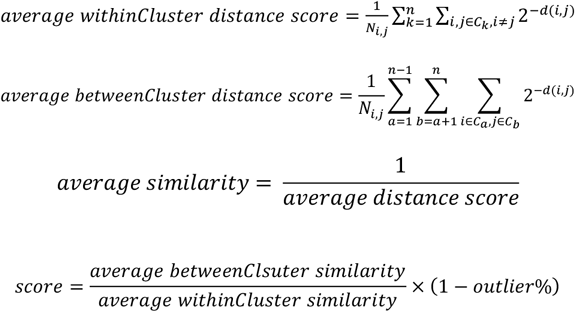

Where d is Euclidean distance between data points and n is the total number of clusters.

The average within-cluster similarity reflects the degree of cohesion and average between-cluster similarity represents the separation of clusters. And the percentage of non-outlier indicate the integrity of information. Obviously, the score becomes bigger when 1. distance among data points in the same cluster is smaller;

1. distance among data points between different clusters is bigger;
2. less outliers are produced. Therefore, the larger the score, the better the clusters are separated.

### 4.5 Eigenvector filtering based on the Greedy strategy

Using PCA, we obtained all the eigenvectors from the standardized data. For each dimension reduction step, we removed one eigenvector in turn and ran the KANN-DBSCAN algorithm with the remaining eigenvectors and computed the corresponding score. Then we reserved the combination of eigenvectors with the highest score for the next dimension reduction. We repeated this process until the dimension reaches 3. In the meantime, if dimensionality reduction fails to divide the data into multiple classes, the algorithm stops. The case with the largest score in all reduction steps is the best, also indicating that the remaining eigenvectors form the optimal PC subspace.

## Acknowledgements

We thank Shengdian Jiang for his comments and suggestions, Zhixi Yun for his help during computational analysis and figure generation. This work has been supported by SEU to support informatics data management and analysis pipeline of full neuronal reconstruction platform. This work was also supported by a MOST (China) Brain Research Project, “Mammalian Whole Brain Mesoscopic Stereotaxic 3D Atlas” (2022ZD0205200 ,2022ZD0205204).

## References

Cajal, S.R.y, & Azoulay, L. (1952) Histologie du systèeme nerveux de l’homme & des vertébrés. Instituto Ramon Y Cajal: Consejo superior de investigaciones cientificas.

Santana, R., McGarry, L. M., Bielza, C., Larrañaga, P., and Yuste, R. (2013) Classification of neocortical interneurons using affinity propagation. Front. Neural Circuits. 7:185. 10.3389/fncir.2013.00185

Zeng, H. (2022) What is a cell type and how to define it? Cell. 185(15), pp. 2739–2755. 10.1016/j.cell.2022.06.031.

Armañanzas, R. and Ascoli, G.A. (2015) Towards the automatic classification of neurons. Trends in Neurosciences. 38(5), pp. 307–318. 10.1016/j.tins.2015.02.004.

Winnubst, J. et al. (2019) Reconstruction of 1,000 projection neurons reveals new cell types and organization of long-range connectivity in the Mouse Brain. Cell. 179(1). 10.1016/j.cell.2019.07.042.

Peng, H. et al. (2015) Bigneuron: Large-scale 3D neuron reconstruction from optical microscopy images. Neuron. 87(2), pp. 252–256. 10.1016/j.neuron.2015.06.036.

Peng, H., Meijering, E. & Ascoli, G.A. (2015) From DIADEM to BigNeuron. Neuroinform. 13, 259–260. 10.1007/s12021-015-9270-9.

Peng, H. et al. (2021) Morphological diversity of single neurons in molecularly defined cell types. Nature. 598(7879), pp. 174–181. 10.1038/s41586-021-03941-1.

Li W, Yan S, Jiang Y, et al. (2019) Research on method of self-adaptive determination of DBSCAN algorithm parameters. Comput. Eng. Appl. 55(5): 1–7.

Scorcioni, R., Polavaram, S. & Ascoli, G. (2008) L-Measure: a web-accessible tool for the analysis, comparison and search of digital reconstructions of neuronal morphologies. Nat Protocols. 3, 866–876. 10.1038/nprot.2008.51.

Jones, A., Overly, C. & Sunkin, S. (2009) The Allen Brain Atlas: 5 years and beyond. Nat Rev Neurosci. 10, 821–828. 10.1038/nrn2722.

Dong, H. W. (2008) The Allen reference atlas: A digital color brain atlas of the C57Bl/6J male mouse. John Wiley & Sons Inc.

Zhang, J., Li, S.Z. and Wang, J. (2005) Manifold learning and applications in recognition. Intelligent Multimedia Processing with Soft Computing. pp. 281–300. 10.1007/3-540-32367-8_13.

Kim, E.J. et al. (2015) Three types of cortical layer 5 neurons that differ in brain-wide connectivity and function. Neuron. 88(6), pp. 1253–1267. 10.1016/j.neuron.2015.11.002.

Gouwens, N.W., Sorensen, S.A., Berg, J. et al. (2019) Classification of electrophysiological and morphological neuron types in the mouse visual cortex. Nat Neurosci. 22, 1182–1195. 10.1038/s41593-019-0417-0.

Cannon, R., Turner, D., Pyapali, G., and Wheal, H. (1998) An online archive of reconstructed hippocampal neurons. Journal of Neuroscience Methods. 84(1-2), pp.49–54.

Ester, M., Kriegel, H.P., Sander, J., Xu, X., et al. (1996) A density-based algorithm for discovering clusters in large spatial databases with noise. Proc. KKD. 96, 226–231.

Gao, L., Liu, S., Gou, L. et al. (2022) Single-neuron projectome of mouse prefrontal cortex. Nat Neurosci. 25, 515–529. 10.1038/s41593-022-01041-5.

Tsiola, A. et al. (2003) Quantitative morphologic classification of layer 5 neurons from mouse primary visual cortex. The Journal of Comparative Neurolog. 461(4), pp. 415–428. 10.1002/cne.10628.

Avrim Blum, Pat Langley. (1997) Selection of relevant features and examples in machine learning. Artif. Intell. 97 (1– 2) (1997), pp. 245–271.

Marx M, Feldmeyer D. (2012) Morphology and physiology of excitatory neurons in layer 6b of the somatosensory rat barrel cortex. Cereb Cortex. 23(12):2803–2817. 10.1093/cercor/bhs254.

DeFelipe J, López-Cruz PL, Benavides-Piccione R, Bielza C, Larrañaga P, Anderson S, Burkhalter A, Cauli B, Fairén A, Feldmeyer D, et al. (2013) New insights into the classification and nomenclature of cortical GABAergic interneurons. Nat Rev Neurosci. 14(3):202–216. 10.1038/nrn3444.

Lida Kanari, Srikanth Ramaswamy, Ying Shi, Sebastien Morand, Julie Meystre, Rodrigo Perin, Marwan Abdellah, Yun Wang, Kathryn Hess, Henry Markram. (2019) Objective Morphological Classification of Neocortical Pyramidal Cells. Cerebral Cortex. Volume 29, Issue 4, Pages 1719–1735. 10.1093/cercor/bhy339.

Lin, X., Zheng, J. (2019) A Neuronal Morphology Classification Approach Based on Locally Cumulative Connected Deep Neural Networks. Appl. Sci. 9, 3876. 10.3390/app9183876.

Deitcher Y, Eyal G, Kanari L, Verhoog MB, Kahou GAA, Mansvelder HD, Kock CPJD, Segev I. (2017) Comprehensive morpho-electrotonic analysis shows 2 distinct classes of L2 and L3 pyramidal neurons in human temporal cortex. Cereb Cortex. 27(11):5398–5414. 10.1093/cercor/bhx226.

Gillette TA, Ascoli GA. (2015) Topological characterization of neuronal arbor morphology via sequence representation: I— motif analysis. BMC Bioinformatics. 16(1):216. 10.1186/s12859-015-0604-2.

Wan Y, Long F, Qu L, Xiao H, Hawrylycz M, Myers EW, Peng H. (2015) BlastNeuron for automated comparison, retrieval and clustering of 3D neuron morphologies. Neuroinformatics. 13(4):487–499. 10.1007/s12021-015-9272-7.

C. M. Procopiuc, M. Jones, P. K. Agarwal, and T. M. Murali. (2002) A Monte Carlo Algorithm for Fast Projective Clustering. In Proc. ACM SIGMOD Int. Conf. on Management of Data (SIGMOD’02). 10.1145/564691.564739.

C. Baumgartner et al. (2004) Subspace selection for clustering high-dimensional data. Fourth IEEE International Conference on Data Mining (ICDM’04). pp. 11–18. 10.1109/ICDM.2004.10112.

Cervantes, E.P., Comin, C.H., Junior, R.M.C. et al. (2019) Morphological Neuron Classification Based on Dendritic Tree Hierarchy. Neuroinformatics. 17, 147–161. 10.1007/s12021-018-9388-7.

Milošević, N.T. et al. (2009) Quantitative analysis of dendritic morphology of the alpha and delta retinal ganglion cells in the rat: A Cell Classification Study. Journal of Theoretical Biology. 259(1), pp. 142–150. 10.1016/j.jtbi.2009.03.011.

Gala, R., Gouwens, N., Yao, Z., Budzillo, A., Penn, O., Tasic, B., Murphy, G., Zeng, H., Sümb ül. (2019) A coupled autoencoder approach for multi-modal analysis of cell types. Advances in Neural Information Processing Systems, 32.

Pfeifer, R., Gómez, G. (2009) Morphological computation-connecting brain, body, and environment. In Creating brainlike intelligence. pp. 66–83. Springer, Berlin, Heidelberg.

Masood, M., Khan, M.N.A. (2015) Clustering techniques in bioinformatics. IJ Modern Education and Computer Science. 1, 38–46. 10.1201/b13091-10.

Karim, M.R. et al. (2021) Deep learning-based clustering approaches for bioinformatics. Briefings in Bioinformatics. 22(1), pp. 393–415. 10.1093/bib/bbz170.

Ascoli, Giorgio A. and Wheeler Diek W. (2016) In search of a periodic table of the neurons: Axonal-dendritic circuity as the organizing principle: Patterns of axons and dendrites within distinct anatomical parcels provide the blueprint for circuit-based neuronal classification. BioEssays, 38(10):969–976. 10.1002/bies.201600067.

